# Arc represses gene expression in IS*605*-family transposons

**DOI:** 10.64898/2026.09.24.754181

**Authors:** Younggi D. Moon, Rimantė Žedaveinytė, Tanner Wiegand, Samuel H. Sternberg

**Affiliations:** Department of Biochemistry and Molecular Biophysics, Columbia University, New York, NY, USA; Howard Hughes Medical Institute, Columbia University, New York, NY, USA

## Abstract

Bacterial insertion sequences (IS) are compact transposable elements that encode proteins required for their mobility and maintenance, yet many also encode accessory proteins with poorly understood functions. For example, IS*605*-family elements often encode a transposase called TnpA and an RNA-guided nuclease called TnpB that supports transposon maintenance, alongside an additional ribbon-helix-helix protein named Arc. Though the roles of TnpA and TnpB have been extensively studied in recent years, the enigmatic function of Arc has not been investigated. Here, we show that Arc acts as a transcriptional repressor to directly bind the transposon’s native promoter sequence regulating TnpA and TnpB gene expression. By systematically testing Arc-containing IS*605* elements, we identified a conserved binding pattern at intergenic transposon sequences neighboring protein-coding genes through chromatin immunoprecipitation and sequencing analyses. We then used fluorescence reporter assays and demonstrated that these intergenic sequences function as strong promoters, and that the presence of Arc dramatically reduces their gene expression. Together, these findings identify Arc as a transposon-encoded transcriptional repressor, revealing a regulatory layer that may promote long-term persistence of IS*605*-family elements by keeping their activity in check. The widespread association of Arc homologs with diverse mobile elements and cellular genes suggests that these compact regulators may more broadly restrain the expression of neighboring genetic machinery across varied genomic contexts.

## INTRODUCTION

Transposable elements (TE) are selfish genetic elements that actively mobilize to new loci in the genome, and are significant drivers of microbial evolution^1–4^. Transposable elements can both benefit and harm the host: they mediate changes that can lead to inactivation of genes^5^, exaptation of transposon genes for host functions such as defense^6–8^, or the transfer of advantageous features such as antibiotic resistance genes^9,10^. However, these elements are parasitic: new integration events might occur in essential host genes, and active transposition can cause genomic instability^11^. Therefore, some mobile elements benefit from less active, tightly regulated transposition^12^. By carefully balancing gene activity, transposons are able to remain proliferative while controlling or suppressing their activity in order to stay ahead of being purged by natural selection.

IS elements are prokaryotic transposons present in over 93% of bacterial genomes, occurring in one or more copies^13^. Insertion sequences (IS) have broadly been defined as bacterial transposons containing only the genes necessary for their mobilization^14^. Yet, despite their definition, a number of IS elements can encode accessory genes beyond transposases^15^. For example, IS*605* elements canonically encode two protein-coding genes, *tnpA* and *tnpB*, only one of which, *tnpA*, encodes a transposase^16^. TnpA is sufficient for mobilization of the transposon to new genomic sites, which happens through a single-stranded DNA intermediate formed during replication, by peel-and-paste mechanism^17–19^. TnpB, on the other hand, is an RNA-guided endonuclease which targets and cleaves the donor site left behind after TnpA excises the transposon from its original location^20,21^. The resulting DNA double-strand break triggers homologous recombination with a newly-synthesized sister chromosome still containing the transposon, thereby restoring the excised copy and preserving the IS*605* element, serving as a transposon retention mechanism^21^ (**Fig. 1a**). Interestingly, recent studies have shown that some IS*605* transposons encode a small putative ribbon-helix-helix (RHH)-family transcription factor, potentially further upending the canonical definition of an IS as encoding only the genes essential for transposition^22^. We henceforth refer to these putative DNA-binding proteins as Arc, in line with previous nomenclature^22^.

**Figure 1.**
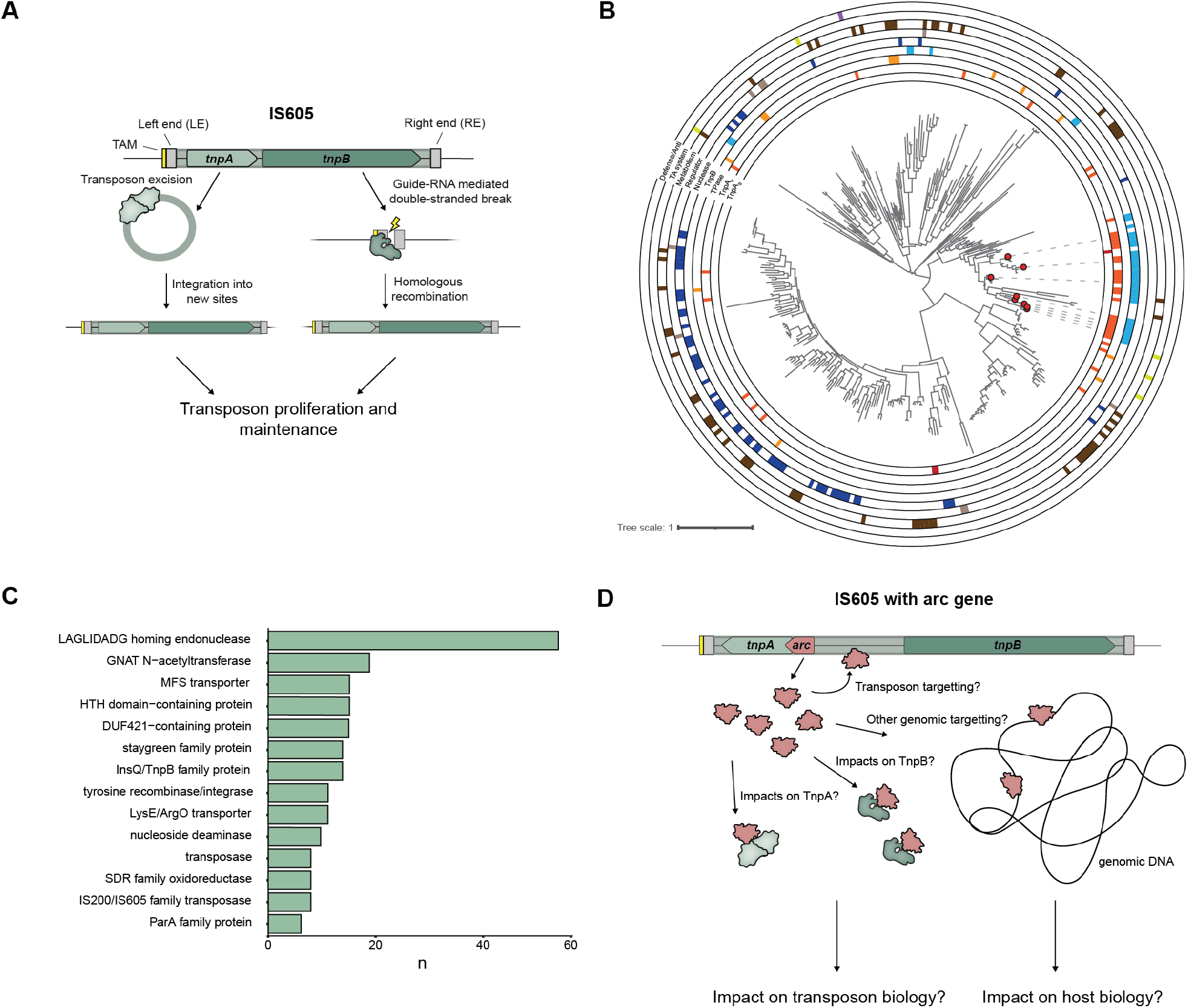
IS605 transposon biology and associations with arc-like RHH repressor genes. **A)** Diagram depicting key roles of TnpA and TnpB in IS605 biology, with TnpA acting as the transposase, mediating transposition events, and TnpB acting as an RNA-guided endonuclease, cleaving regions in the genome in which the transposon was excised, leading to homologous recombination and maintenance of the transposon at the original site. **B)** Phylogenetic tree displaying all cluster members for a clade of Arc homologs associated with transposon proteins including IS605 native TnpA and TnpB. Additional annotations are labeled. Sequences representing experimental candidates are highlighted with red circles. **C)** Genes located immediately upstream or downstream of *arc,* within the Arc-encoding loci of the clade shown in Fig. 1b. In addition to transposon constituents (e.g., transposases, recombinases, TnpB nucleases), homing endonucleases and metabolic components are frequently associated. **D)** Example of an IS605 element containing an *arc* gene. This schematic highlights the potential regulatory role that Arc may play, possibly binding to either the transposon, other genomic regions, or even possibly interacting with transposon genes such as TnpA and TnpB.

IS*605*-encoded Arcs are a part of a broader class of RHH-family proteins, which have been shown to have diverse roles: MetJ acts as a transcriptional repressor in methionine biosynthesis, and ω, which represses the expression of key genes involved in controlling plasmid copy number and segregation^23–25^. Some RHH proteins have been shown to act as transcriptional activators, such as AlgZ, which activates gene expression for an alginate synthesis pathway in *Pseudomonas aeruginosa*^26–28^. RHH-domain containing proteins can also be found in viral genomes, such as aCcr1 which allows the virus to disturb the cell cycle to its own advantage^29^. RHH homologs also act in autoregulatory mechanisms within toxin-antitoxin systems which allow for the RHH to also control expression of toxin-antitoxin genes as well as its own expression^30,31^. These regulators are widespread and play varied roles in both mobile genetic parasites as well as in the host organisms themselves, suggesting that IS*605* encoded Arc may also play a regulatory role.

IS elements often control their own gene expression through diverse mechanisms. Some IS elements depend on selective pressure to mediate bursts in transposase gene expression, as is the case for IS256 in pathogenetic *E. faecium* or *E. faecalis* when facing antibiotic or phage-mediated selection respectively^32^. In other cases, IS elements harbor transient promoters that are only active under the formation of a transposition intermediate circular junction that connects the transposon ends, temporarily activating the promoter and therefore also transposon gene expression^33,34^. Some IS elements simply rely on weak endogenous promoters to keep transposase levels at a lower constitutive level^35–37^. All these mechanisms act as expressional tuning which allows for particular conditions in which transposon genes are expressed higher to promote transposition and transposon proliferation. This type of gene regulation appears to be a prevalent feature of IS element biology, and the presence of *arc* genes encoded within some IS*605* elements presents the possibility of IS-encoded gene regulation.

Here we show that Arc encoded within IS*605*-family transposons bind double-stranded DNA in a sequence-specific manner. The sequence motif that Arc binds overlays the predicted promoter governing the expression of adjacent IS*605* transposon encoded genes, including the *arc* gene itself. We then show DNA binding leads to transcriptional silencing of downstream expression, revealing a new mechanism of transposon gene regulation by the Arc repressor, and offering insight into how these elements adapt over time to be safely maintained in the genome.

## RESULTS

### Arc proteins in IS*605*-family transposons

The presence of *arc* genes within the boundaries of some IS*605*-family elements suggested a likely conserved role in the transposon life cycle^22^. To explore this possibility, we identified a large family of Arc homologs through both sequence and structural homology utilizing BLASTp, PSI-BLAST, and Foldseek^38–40^. The surrounding genomic neighborhoods and adjacent genes were then extracted to look for potential transposon components or other associated genes. This approach identified 8,024 unique Arc-like proteins, and while some homologs were indeed positioned near transposon-encoded proteins (i.e., transposases and TnpB-family nucleases), the majority of Arc-like proteins are encoded immediately adjacent to functionally diverse genes — including toxin-antitoxin systems, metabolic effectors, and other Arc-family DNA-binding proteins (**Fig. S1**).

Among these diverse associations, we identified a clade of Arc proteins with numerous transposon-associated genes, including tyrosine recombinases/integrases and other transposase genes (**Fig. 1b**). A monophyletic clade of Arc proteins encoded both TnpA and TnpB immediately proximal to *arc*, suggesting these Arc proteins occur within IS elements. Many of these systems feature a *tnpB* gene encoded in one direction and *tnpA* and *arc* genes encoded in the other direction, with opposite strandedness, suggesting that the intergenic space possibly acts as a bidirectional promoter region for expression of transposon genes. Notably, within this subgroup, Arc proteins were also found adjacent to homing endonuclease and acetyltransferase genes, suggesting additional functional associations among closely related Arc proteins (**Fig. 1c**).

To investigate Arc-containing IS*605* elements further, we selected systems with *arc* genes encoded in close proximity to either *tnpB* genes or *tnpA* genes, and then tested their functionality in a heterologous *E. coli* host. First, to survey these transposons for TnpA catalyzed DNA excision, we utilized a PCR-based excision assay. By co-transforming cells with a mini-IS element and TnpA overexpression plasmid, we were able to detect transposon excision via PCR and gel electrophoresis analysis (**Fig. S2a**); sequencing analyses further confirmed that shorter-mobility bands indeed corresponded to faithful excision products of the mini-IS*605* elements (**Fig. S2b**). Some of these IS*605* elements also exist in multiple copies in their genomes, which together with the excision data indicates these transposons are likely active for transposition. We next assessed the RNA-guided DNA cleavage activity of transposon-encoded TnpB proteins, which can promote transposon maintenance and spread^21,22,41^; some TnpB homologs lack nuclease activity but retain RNA-guided DNA binding and have evolved roles as transcriptional repressors^42^. To determine whether the TnpB proteins associated with Arc-containing elements retain DNA cleavage activity, we tested them using a plasmid interference assay (**Fig. S3a**). TnpB cleavage activity varied across the tested homologs (**Fig. S3b**), with substitutions in the catalytic DDE motif providing a possible explanation for the lack of detectable activity in some cases (**Fig. S3c**). Together with the TnpA excision assays, these results demonstrate characteristic IS*605*-family activities in a subset of the Arc-containing elements tested.

We next investigated how Arc might contribute to the life cycle of these elements. Given the established DNA-binding activity of related RHH-family proteins^25^, we hypothesized that Arc might bind the transposon itself or other genomic sequences to regulate gene expression. Arc has also been proposed to interact with the TnpB–ωRNA complex^43^, suggesting a possible role in modulating TnpB function. These models provided a starting point for investigating Arc activity in IS*605*-family elements (**Fig. 1d**).

### Arc binds IS*605*-family transposon promoters

Since RHH homologs tend to bind to promoters, we searched for promoters responsible for the expression of the transposon native *arc*, *tnpA*, and *tnpB* genes. Utilizing the previously identified homologs, their associated genes, and the promoter prediction software BPROM^44^, we identified predicted promoter motifs in the intergenic region between the transposon genes (**Fig. S4a**). In many cases we identified two promoter motifs with promoter driving expression of *tnpB* and a second promoter in the opposing direction likely regulating the expression of *tnpA* and *arc* (**Fig.S4b**).

To test for the sequence-specific binding activity of IS*605*-associated Arc homologs, we performed chromatin immunoprecipitation sequencing (ChIP-seq) assays with Arc homologs that exhibit IS*605*-family transposon association. We tagged the C-terminus of the Arc protein with 3x-FLAG tag and overexpressed it heterologously in BL21(DE3) *E. coli* cells in the presence of a target plasmid (pTarget) encoding both the native *tnpB* gene sequence and the bi-directional transposon promoter. Arc-bound DNA was then immunoprecipitated and sequenced to determine Arc binding sites (**Fig. 2a**). By testing Arc from two autonomous IS*605* elements native to *Petroclostridium xylanilyticum*^45^ and *Clostridium botulinum*, we observed a substantial enrichment of reads mapping to the transposon’s putative promoter, identifying this region as the predominant Arc binding site (**Fig. 2b**). Both homologs also produced weaker peaks at sites in the *E. coli* BL21(DE3) genome with partial sequence similarity to the native transposon target (**Fig. 2c**).

**Figure 2.**
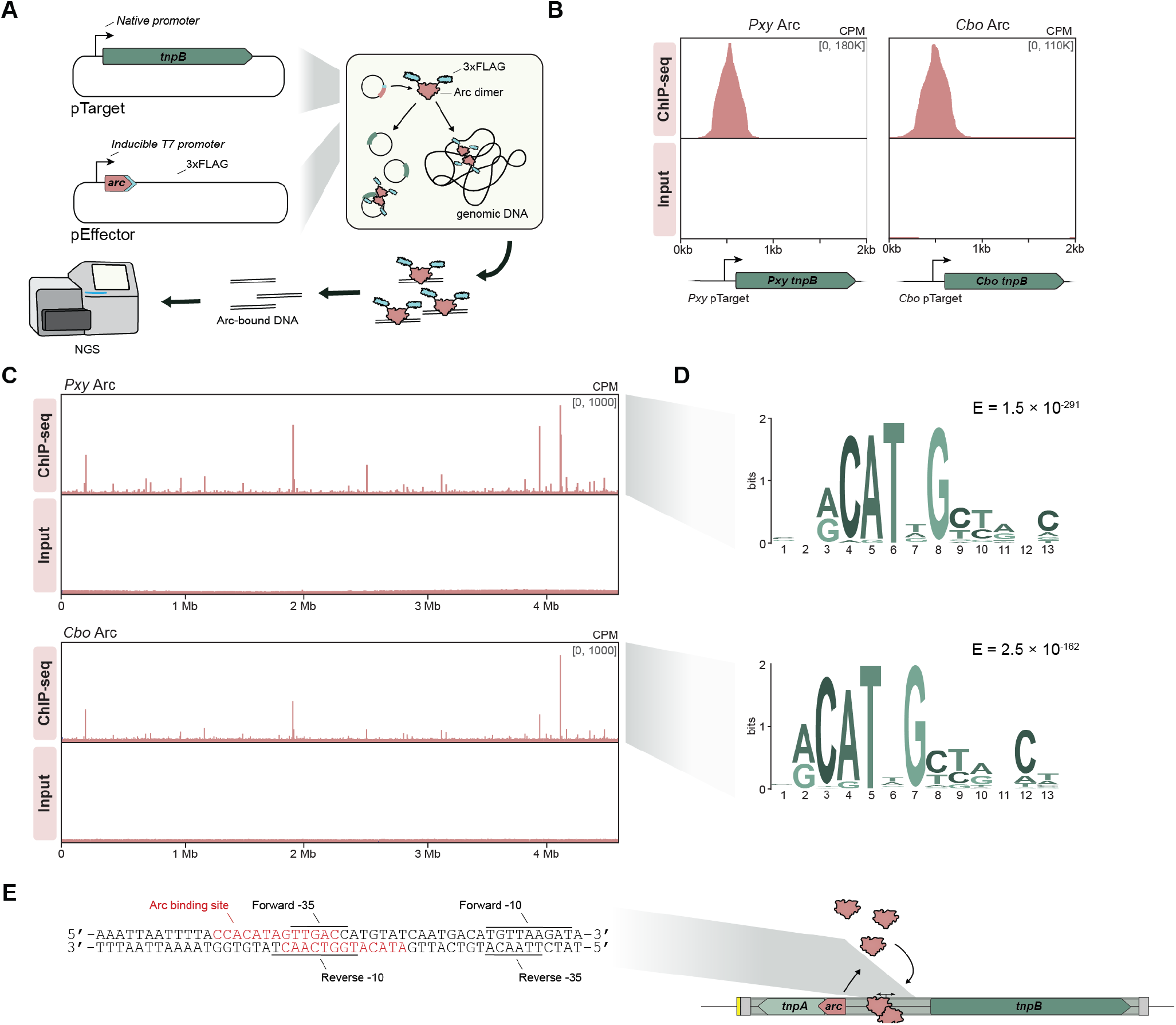
IS605-encoded RHH repressors bind to putative promoters driving transposon gene expression. **A)** Experimental workflow for ChIP-seq assay. One vector encoding the transposon’s native promoter and TnpB gene (pTarget) and a second vector with IPTG inducible overexpression of a 3xFlag tagged native Arc protein (pEffector) were both transformed and cultured for subsequent ChIP-seq analysis allowing for determination of binding sites through enrichment of reads over binding sites. **B)** ChIP-seq data for experiments done with *C. botulinum* and *P. xylanilyticum* derived transposons. This data shows reads that map to the pTarget vector, and both show strong enrichment of reads directly above the putative native transposon promoter, indicating that the 3xFlag tagged arc-like RHH repressors are binding specifically to this DNA sequence. **C)** ChIP-seq data mapped to the genome of BL21(DE3) *E. coli*. Enrichment of reads over any given peak is much less than the enrichment over the transposon’s native promoter sequence. This indicates that over-expression may induce some off-target binding for partial binding sites. **D)** Binding motifs for arc-like RHH repressors for *P. xy* and *C. bo* derived from the genomic peaks using MEME-ChIP to determine consensus sequences, which are near identical for both *P. xy* and *C. bo*. E, E-value significance. **E)** Labeling of *P. xy* consensus motif on the native sequence that contains putative promoters (predicted by BProm) for both *tnpB* as well as the *arc* and *tnpA*. Labeled in red are the motifs derived from genomic peaks. *P. xy* has binding motifs positioned over the −35 motif corresponding to the *tnpB* promoter and the −10 motif corresponding to the *tnpA* and *arc* promoter.

Motif analysis of genomic ChIP-seq peaks identified with MACS3 revealed nearly identical consensus binding sequences for the two Arc homologs (**Fig. 2d**). Mapping these motifs onto the native transposon sequences placed Arc binding sites within the predicted promoter regions. In the *P. xylanilyticum* element, oppositely oriented Arc binding motifs overlap the predicted −35 and −10 elements of the divergent promoters (**Fig. 2e**). This arrangement is consistent with a model in which two Arc dimers bind the promoter region, as observed for other RHH-family regulators^25^. We extended these analyses to additional Arc homologs from IS*605*-family elements encoding either *arc* and *tnpB* alone, or all three genes, *arc*, *tnpA*, and *tnpB* (**Fig. S5, S6**). Across both groups, several homologs showed strong enrichment at the native transposon sequence, accompanied by weaker peaks in the *E. coli* genome (**Fig. S5a,b, S6a,b**). Motifs derived from these genomic peaks also mapped to the native transposon sequence (**Fig. S5c, S6c**). Other homologs showed no detectable sequence-specific binding to either the transposon or the *E. coli* genome under the conditions tested. Thus, although binding activity varied among homologs, the native transposon sequence was consistently the predominant target whenever sequence-specific binding was detected.

Together, these results show that multiple Arc homologs recognize specific DNA sequences within their associated IS*605*-family elements. The overlap of Arc binding sites with predicted promoter elements supports a role in regulating transposon gene expression.

### Arc binding mediates transcriptional repression

The overlap of Arc binding sites with the native transposon promoter, together with the established repressive activity of many RHH-family proteins, suggested that Arc might repress transposon gene expression. We therefore developed a fluorescence reporter assay to test this hypothesis.

We co-transformed *E. coli* with an inducible Arc expression plasmid and a reporter plasmid expressing RFP from the native *tnpB* promoter. Reporter fluorescence was normalized to OD_600_ to account for differences in cell density (**Fig. 3a**). To assess the activity of this heterologous promoter in *E. coli*, we compared it with a strong constitutive *E. coli* promoter and a random non-promoter sequence as positive and negative controls, respectively (**Fig. 3b**). In the absence of Arc, the transposon and constitutive promoters drove similar levels of RFP fluorescence, whereas the non-promoter control produced essentially no RFP signal. Arc expression substantially reduced fluorescence from the transposon promoter but had no detectable effect on the constitutive promoter, supporting promoter-specific repression (**Fig. 3c**). Time-course measurements in liquid culture further showed that normalized fluorescence eventually significantly declined in Arc-expressing cells while continuing to increase in cells lacking Arc (**Fig. 3d**). Together with the ChIP-seq results, these findings support a model in which Arc binds the native transposon promoter to repress downstream gene expression.

**Figure 3.**
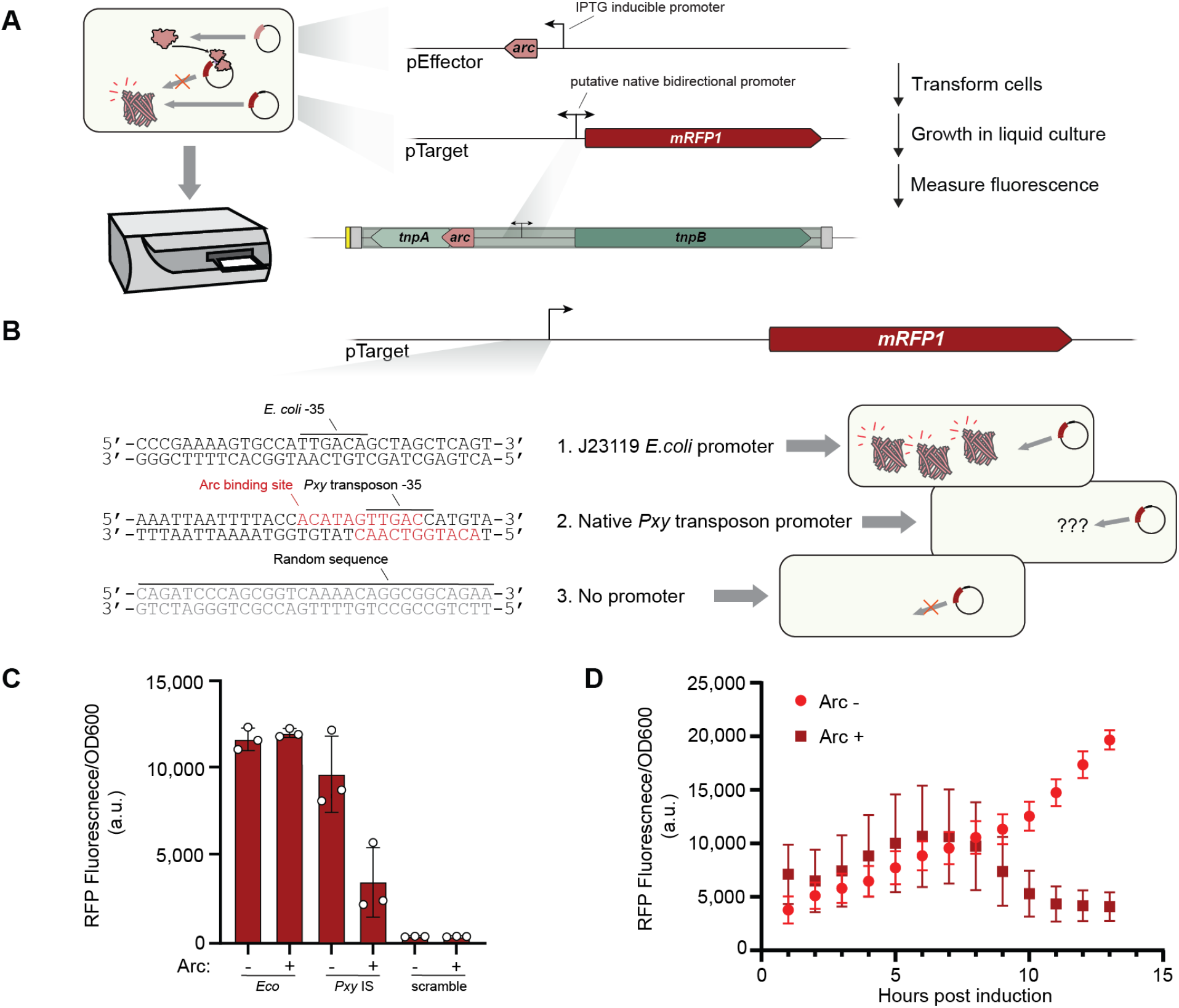
Binding of arc-like RHH-repressors to native promoters reduces downstream protein expression. **A)** Schematic of the fluorescence reporter assay. *E. coli* BL21(DE3) cells were co-transformed with an inducible Arc expression vector (pEffector) and a reporter plasmid (pTarget) expressing mRFP1 from the native transposon promoter. Reporter fluorescence was normalized to OD_600_. **B)** Promoter sequences used in the reporter constructs, including the constitutive *E. coli* J23119 promoter, the native *P. xylanilyticum* transposon *tnpB* promoter, and a scrambled sequence serving as a negative control. Predicted promoter elements and the Arc binding site are indicated. **C)** Endpoint measurements of normalized RFP fluorescence for the three reporter constructs in **b**, with Arc expression (+) or an empty-vector control (–). All conditions contained 0.5 mM IPTG. Arc expression was associated with lower mean fluorescence from the native transposon promoter, whereas fluorescence from the constitutive promoter was similar with and without Arc. **D)** Time course of normalized RFP fluorescence from the native transposon promoter with Arc expression or an empty-vector control. The trajectories diverged approximately 8 h after induction, with fluorescence continuing to increase in the control and declining in Arc-expressing cells. Data in **C** and **D** represent means with error bars indicating SD from three biological replicates.

### DNA recognition underlies Arc-mediated repression

Structural modeling with AlphaFold 3 and visualization in ChimeraX suggested that Arc adopts a canonical RHH architecture, with two β-strands forming a DNA-binding surface positioned in the major groove of DNA^46,47^ (**Fig. 4a**). The model identified Asn5, Asn7, and Thr9 as candidate DNA-contacting residues, and an alignment of nine closely related Arc homologs revealed conserved positions within both the helical regions and the β-strands, consistent with proposed roles in dimerization and DNA recognition, respectively^25,48^ (**Fig. 4b**).

**Figure 4.**
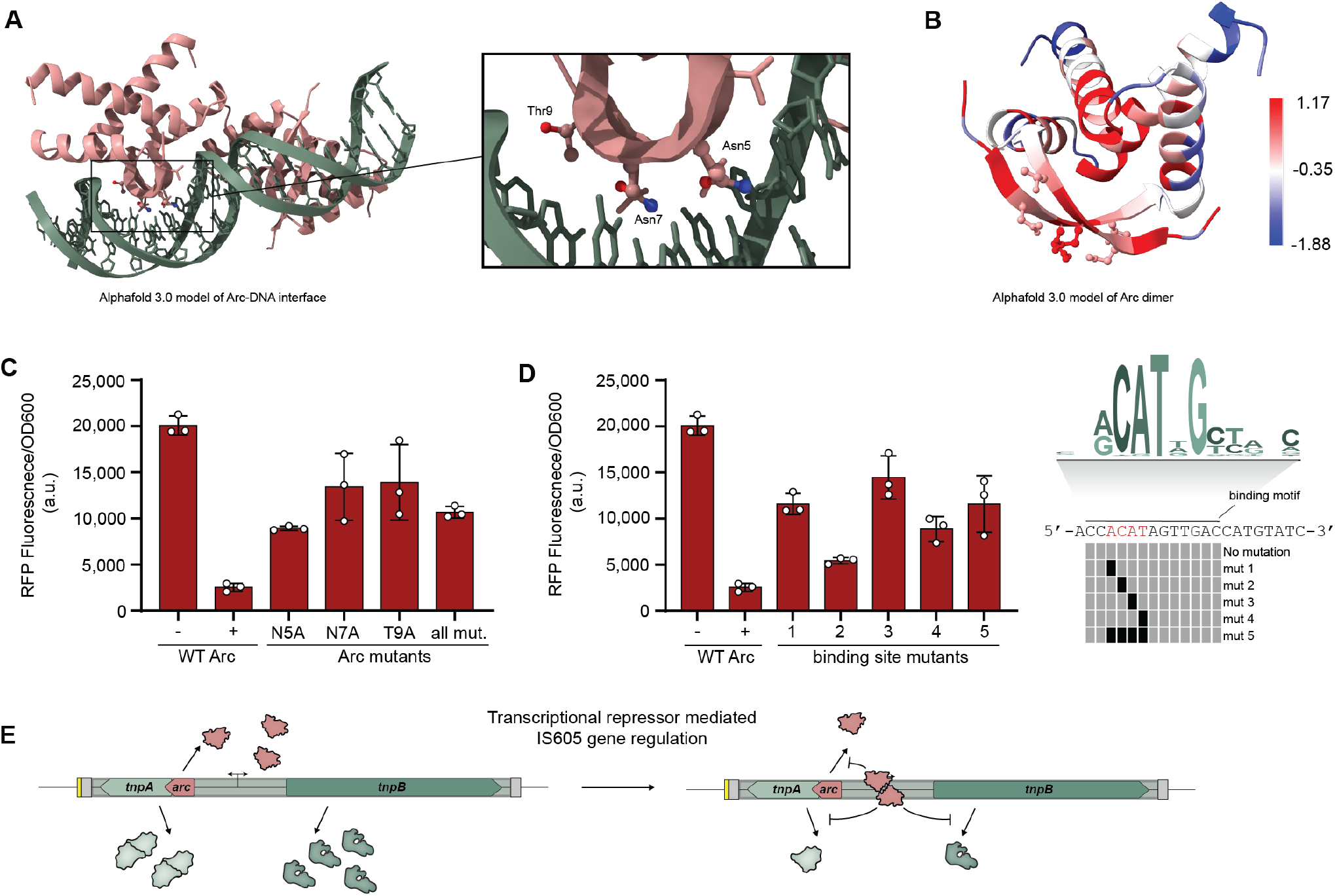
Mutational perturbations show RHH binding is necessary for downstream impacts on transposon native promoter. **A)** AlphaFold 3 model of two Arc dimers bound to promoter DNA, visualized in ChimeraX. The inset highlights predicted DNA-contact residues Asn5, Asn7, and Thr9. **B)** Sequence conservation mapped onto the predicted Arc dimer structure, based on an alignment of nine closely related homologs. Colors indicate AL2CO conservation scores^48^, from low (blue) to high (red). **C)** Effects of alanine substitutions at predicted DNA-contact residues on Arc-mediated repression of an RFP reporter. “All mut.” denotes the N5A/N7A/T9A triple mutant. Controls contain an empty vector (−) or wild-type Arc (+). **D)** Effects of binding-site substitutions on reporter repression by wild-type Arc. Substitutions shown at right replace the indicated nucleotides with their complementary bases while preserving the −35 promoter element. Wild-type controls are as in **C**. Data in **C** and **D** represent means with error bars indicating SD from three biological replicates. **E)** Proposed model of Arc-mediated transcriptional repression in IS605-family elements. Arc binding represses expression from the *tnpB* promoter; repression of *tnpA* and autoregulation of *arc* remain proposed extensions of this model.

To test the contribution of the predicted DNA-contacting residues to repression, we substituted Asn5, Asn7, and Thr9 with alanine, individually or in combination, and measured RFP reporter fluorescence. All variants showed significantly reduced repression relative to wild-type Arc (**Fig. 4c**), supporting a functional role for the predicted DNA-binding surface in Arc-mediated transcriptional control. We next tested the contribution of the cognate DNA sequence by introducing substitutions into the Arc binding motif while leaving the −35 promoter element intact. Each tested substitution significantly reduced Arc-mediated repression, including single-nucleotide changes (**Fig. 4d**). None fully restored reporter fluorescence to the level observed without Arc, potentially reflecting residual recognition of the partially intact binding site.

Together, the structural predictions and mutational analyses support a model in which Arc recognizes a specific site within the native transposon promoter through its RHH DNA-binding surface and represses downstream gene expression (**Fig. 4e**).

## DISCUSSION

Insertion sequences are ubiquitous in prokaryotic genomes, and understanding how they interact with their hosts could help explain how IS elements adapt to host genomes. Our findings reveal the role of Arc, a conserved accessory protein encoded within some IS*605*-family elements. We show that Arc binds the native transposon promoter, thereby restricting expression driven by the promoter of *tnpB*, which encodes an RNA-guided endonuclease involved in transposon retention^21^. Structural predictions and mutational analyses further support a canonical RHH mechanism of DNA recognition underlying this repression (**Fig. 4e**). Given the overlap of Arc binding sites with two predicted divergent promoters, Arc may also repress its own expression and that of *tnpA*, potentially coordinating expression of multiple transposon genes simultaneously. Building on these findings in heterologous *E. coli* assays, studies in native hosts could establish how Arc-mediated repression responds to physiological conditions and whether its relief permits increased transposition. Overall, these findings reveal a regulatory mechanism that may promote long-term maintenance of IS*605*-family elements in the host genome by carefully controlling their gene expression.

Arc-mediated repression adds to a growing set of examples in which transposon-encoded regulatory proteins control element activity. In Type V-K CRISPR-associated transposons (CAST), CvkR directly represses expression of *cas12k* and *tnsB* through a MerR-family winged helix-turn-helix domain, structurally distinct from the RHH fold of Arc^49^. Interestingly, other Type V-K systems, including the recently described V2 subgroup, encode predicted Arc-like RHH regulators, suggesting that related proteins may regulate gene expression across distinct transposon families^50^. In IS*91*, the accessory protein Orf121 inhibits transposition, and its expression is associated with reduced *tnpA* transcript levels^51^. The overlapping *orf121-tnpA* gene arrangement also influences transposase expression, raising the possibility that the analogous overlap between *arc* and *tnpA* contributes an additional layer of regulation in IS*605*-family elements. Together, these examples demonstrate highly diverse mechanisms through which transposons may coordinate gene expression and restrain their own activity.

Our findings also highlight the regulatory potential of small proteins that can escape standard genome annotation. Some Arc homologs examined here fall within the ≤50-amino-acid class of micro-proteins^52–54^. Comparative sequence analysis also identified *arc* genes that were absent from existing annotations. Together with the discovery of numerous phage-associated microproteins^55^, these observations suggest that small, overlooked proteins may contribute broadly to the regulation of mobile gene elements.

IS*605*-family transposons have given rise to diverse molecular systems relevant to biotechnology and human health. Our work expands this functional repertoire by identifying Arc-mediated transcriptional repression as a mechanism for controlling transposon gene expression. Although insertion sequences were initially defined by their apparent simplicity, our work reveals their surprising complexity. Further study may uncover new biological systems with biotechnological potential and clarify how IS elements persist over evolutionary time.

## METHODS

### Reagents

All necessary reagents and specialized reagents for experiments are listed in the methods below alongside their manufacturer.

### Biological Resources

All Arcs tested, strains, plasmids, and oligonucleotides used for this study are listed in **Supplementary Table 2-5** with full sequences and corresponding identification numbers.

### Statistical Analyses

All analyses were performed on GraphPad Prism 11.1.0. All statistically tested comparisons involved three separate biological replicates (separate bacterial colonies) per sample group. A “significant” reduction or difference was stated if an unpaired Welch’s t-test was performed with a resulting p ≤ 0.05 (two-sided). In cases where biological replicates had technical replicates, technical replicates were averaged prior to analysis of 3 individual biological replicates.

### Bioinformatic analyses of Arc proteins

To construct a dataset of Arc protein sequences, tandem sequence and structure-based homology searches were undertaken. A previously described set of 770 Arc sequences^22^ were used to query a local version of the NR database (downloaded on April 16, 2025) in a PSI-BLAST^56^ search [E-value < 0.1] run to convergence, resulting in 849 unique Arc sequences. Concurrently, an Arc homolog from *Petroclostridium xylanilyticum* (WP_094548610.1) was queried against the NR database in a BLASTp search. The structures of eight resultant hits were then predicted with AlphaFold 3^47^, and these predicted structures were used to query Foldseek^39^. The resulting 2,812 Arc-like proteins from the Foldseek search were then searched against the local NR database, retrieving 19,407 Arc proteins. Arc protein hits from the PSI-BLAST approach and Foldseek pipeline were then concatenated and de-duplicated, and the locus encoding each protein (upstream gene + *arc* + downstream gene) was retrieved for each unique Arc homolog. This final dataset consists of 10,374 unique Arc proteins with an accompanying locus.

Arc-RHH domains are sometimes embedded within larger polypeptides. To separate out stand-alone Arc regulators, Arc homologs that spanned less than 200 amino acids were extracted from the full dataset, resulting in a dataset of 8,024 unique Arc proteins. To assess which genes are stably associated with *arc*, translated sequences of open reading frames (ORFs) flanking *arc* were scanned with a suite of profile hidden Markov models (HMMs)^57^. First — to assess the presence of transposon-encoded proteins — Arc-neighboring ORFs were scanned with Pfam^58^ HMMs built from TnpB (PF05717.18 and PF01385.24), a Pfam HMM built from IS607 TnpA (PF00239.26), HMMs from ISEscan^59^, and Pfam HMMs in the RNAse_H clan (CL0219) which includes numerous transposases. Arc-neighboring ORFs were scanned against all of the HMM sets described above (TnpB, IS607 TnpA, ISEscan, and the RNase_H clan). Where an ORF received hits from more than one resource, the annotation was assigned according to the priority order listed above (TnpB > TnpA > ISEscan > RNase_H clan), with the higher-priority resource’s annotation retained. The remaining ORFs, which did not receive an annotation from those collected HMMs, were annotated with eggNOG-mapper^60^.

To construct an approximate phylogenetic tree of all Arc sequences, the dataset of 8,024 unique stand-alone Arc proteins was clustered with MMseqs2^61^ [easy-linclust --min-seq-id 0.6 --min-aln-len 50 --cov-mode 5]. 2,175 Arc cluster representatives were then aligned with MAFFT^62^ [EINSI setting], and the resulting alignment was trimmed with Trimal^63^ to remove columns composed of >90% gaps. The trimmed alignment was then used to construct a phylogenetic tree with FastTree^64^ (-wag -gamma). This tree of cluster representatives (shown in **Fig. S1**), is an approximate ML tree used to order representatives for visualization, and deep relationships should not be interpreted due to the high sequence divergence across this dataset of Arc homologs.

To construct a more refined tree of transposon-associated Arc homologs (**Fig. 1b**), a clade of cluster representatives that encompassed experimental candidates and their neighboring branches was extracted from the **Fig. S1** tree. All of the cluster members represented by these tips (n = 362 unique Arc homologs) were then aligned with MAFFT (EINSI), and a tree was built with FastTree (-wag -gamma).

### Cloning and strain generation

All transposon sequences and genes including TnpA, TnpB, and Arc were ordered from and synthesized by Twist Biosciences. Codon-optimization was done for all transposon genes as necessary but excluded any TnpB protein-coding sequences within 260-bp of the predicted right-end of the IS*605* element as to prevent interference with a potential TnpB guide-RNA scaffold. ChIP-seq pTarget and pEffector plasmids were both synthesized utilizing Gibson assembly. pTarget synthesis involved cloning the native transposon sequence into pCDFDuet-1 vectors with the final construct capturing both the putative promoter sequence and the *tnpB* gene. pEffector plasmids were cloned by encoding *arc* genes downstream of T7 promoter into pCOLADuet-1 vectors. 3xFlag-tags on Arc and TnpB proteins were introduced via around-the horn PCR using primers with overhangs. RFP expression plasmids were either cloned utilizing around-the-horn PCR by inserting the putative transposon promoter upstream of an RFP into a previously existing RFP-expression vector present in the lab (pSL5895) or synthesized by Genscript utilizing a vector that had already been cloned using the prior method (pSL9245). Any subsequent variants of the plasmids listed were cloned utilizing Gibson assembly, around-the-horn PCR, and Golden Gate Assembly, with all PCR steps utilizing Q5 polymerase. Plasmids cloned were used to transform NEB Turbo competent cells (New England Biolabs) and purified using QIAprep Spin Miniprep Kit (Qiagen). Sequence verification was performed with Sanger sequencing through Genewiz or Plasmidsaurus Whole Plasmid Sequencing.

### Chromatin immunoprecipitation and sequencing (ChIP-seq)

*E. coli* BL21(DE3) cells were transformed with pTarget, encoding the transposon’s native promoter, and pEffector with an inducible arc-3x-FLAG gene. The cells were plated onto LB agar plates (200 µg mL^−1^ spectinomycin, 50 µg mL^−1^ kanamycin). Single colonies were picked from each plate and grown overnight in 20 mL of LB supplemented with 200 µg mL^−1^ spectinomycin, 50 µg mL^−1^ kanamycin and 0.5 mM IPTG. ChIP-seq was done following the previously established protocol^65^. In brief, to each of the liquid cultures, 20 mL of LB was added. For cross-linking Formaldehyde (Thermo Fisher Scientific) was added to a concentration of 0.90% (w/v), and the solution was mixed by vortexing, followed by the nutation of each tube for 20 minutes at room temperature. After this, 4.6 mL of 2.5 M glycine was added to stop cross-linking and the tubes were once again nutated for 10 minutes at room temperature. Following this cells were pelleted in a centrifuge at 4000 x*g* for 10 minutes at 4 °C. All buffers for the following were sterile filtered using a 0.22 µL filter and all following steps were performed on ice. Supernatant was discarded and the pellets resuspended in 40 mL of TBS buffer (200 mM Tris–HCl, pH 7.5 at 4 °C, 1.5 M NaCl). The OD_600_ was measured and all samples were standardized by transferring the equivalent of 40 mL of OD_600_ = 0.6 into another set of tubes. These samples were then again centrifuged at 4000 x*g* for 10 minutes at 4 °C. The supernatant was removed and the cells were frozen with liquid nitrogen and stored at −80 °C for future steps

A 5 mg mL^−1^ BSA solution was made by dissolving bovine serum albumin (GoldBio) in 1 PBS buffer (Gibco). For immunoprecipitation, 25 µL of Dynabeads Protein G (Thermo Fisher Scientific) per sample were initially all aliquoted into a single tube, and washed with 1 mL of BSA solution three times at room temperature and resuspended in BSA solution in the initial bead volume. Monoclonal anti-Flag M2 antibodies (Sigma-Aldrich) were added (4 µL per sample) followed by rotation at 4 °C for 3 hours to conjugate the antibodies to the beads. Separately, the previously flash-frozen cell pellets were thawed and resuspended in FA lysis buffer 150 (50 mM HEPES-KOH pH 7.5, 0.1% (w/v) sodium deoxycholate, 0.1% (w/v) SDS, 1 mM EDTA, 1% (v/v) Triton X-100, 150 mM NaCl) with protease inhibitor cocktail (Sigma-Aldrich). These samples were then transferred to a Covaris 1 mL milliTUBE AFA Fiber and then sonicated either on a Covaris M220 Focused-ultrasonicator or on a Covaris LE220 Focused-ultrasonicator. The M220 had the following settings: a minimum temperature of 4 °C, set point of 6 °C and a maximum temperature of 8 °C, peak power of 75, a duty factor of 10, Cycles/Burst of 200, and a sonication time of 17.5 min. The LE220 used the following settings: a minimum temperature of 4 °C, set point of 6 °C and a maximum temperature of 8 °C, peak power of 420, a duty factor of 30, Cycles/Burst of 200, and a sonication time of 17.5 min as well. Samples were then centrifuged at 20,000 x*g* at 4 °C for 20 minutes and the supernatant was transferred to a fresh tube and the pellet was discarded. 10 µL of the supernatant were transferred to a separate tube, flash frozen in liquid nitrogen, and kept at −80 °C as non-immunoprecipitated control.

The beads conjugated with antibodies were washed four times using BSA solution at 4 °C before being resuspended in FA Lysis Buffer 150 with protease inhibitor (30 µL per sample). To each sonicated sample, 31.5 µL of resuspended beads were added, and the samples were rotated for 16 hours at 4 °C. After overnight incubation, the beads were washed in accordance with previous papers^66^. Six total washes were done with the following buffers: (1) two washes with FA lysis buffer 150 (without protease inhibitor); (2) one wash with FA lysis buffer 500 (50 mM HEPES-KOH pH 7.5, 0.1% (w/v) sodium deoxycholate, 0.1% (w/v) SDS, 1 mM EDTA, 1% (v/v) Triton X-100, 500 mM NaCl); (3) one wash with ChIP wash buffer (10 mM Tris-HCl pH 8.0, 250 mM LiCl, 0.5% (w/v) sodium deoxycholate, 0.1% (w/v) SDS, 1 mM EDTA, 1% (v/v) Triton X-100, 500 mM NaCl); and (4) two washes with TE buffer 10/1 (10 mM Tris-HCl pH 8.0, 1 mM EDTA). After the last wash the beads were resuspended in 200 µL of fresh ChIP elution buffer (1% (w/v) SDS, 0.1 M NaHCO_3_) and suspensions were incubated at 65 °C for 1.25 hours while vortexing every 15 minutes in order to keep the beads suspended in the buffer. During this time, the previously aliquoted non-immunoprecipitated input control samples were thawed and 190 µL of ChIP elution buffer was added to each tube. Subsequently 10 µL of 5 M NaCl was also added. After incubation, the immunoprecipitated samples were put onto the magnetic rack and the supernatant was put into a new tube and combined with 9.75 µL of 5 M NaCl. All samples including input control samples were incubated at 65 °C overnight to reverse cross-link. The following day, 1 µL of 10 mg mL^−1^ RNase A (Thermo Fisher Scientific) was added to each sample and incubated for 1 hour at 37 °C. Then, 2.8 µL of 20 mg mL^−1^ proteinase K (Thermo Fisher Scientific) and 1 hour of incubation at 55 °C. Samples were then column purified using QIAquick spin columns (QIAGEN) and eluted in 40 µL TE buffer 10/0.1 (10 mM Tris-HCl pH 8.0, 0.1 mM EDTA).

Sample concentrations for both immunoprecipitated and non-immunoprecipitated input samples were quantified using the DeNovix dsDNA Ultra High Sensitivity Kit. DNA amounts were standardized across all input and immunoprecipitated samples for Illumina libraries prepared using the NEBNext Ultra II DNA Library Prep for Illumina (NEB). After adapter ligation, adapter ligated DNA was cleaned up utilizing AMPure XP beads. AMPure XP beads were allowed to warm to room temperature and resuspended. Then to each adapter-ligated DNA sample 87 µL (0.9 sample volume) of AMPure beads were added and gently vortexed. Samples were incubated at room temperature for 5 minutes before samples were placed on a magnetic rack. Once the beads had settled, the supernatant was removed and discarded. Beads were washed with 200 µL of freshly made 80% ethanol three times. The beads were then allowed to air dry for up to 8 minutes before tubes were removed from the magnetic rack. DNA was then eluted from the beads by adding 17 µL of TE buffer 10/0.1 and vortexing. These were then incubated for 5 minutes at room temperature and the supernatant was kept for PCR amplification while the beads were discarded. PCR amplification for 12 cycles was done to add barcodes. Then, the sample volume was brought up to 50 µL by adding TE Buffer 10/0.1. Then, for two sided size selection, 27.5 µL (0.55 sample volume) of AMPure XP beads (Beckman Coulter) were added to the DNA and the sample placed onto a magnetic rack. The supernatant was then kept with the beads being discarded. Following this, 17.5 µL (0.35 sample volume) of the AMPure beads were added to the DNA. Once again the sample was placed onto a magnetic rack. The supernatant was discarded and the beads were kept, and washed with 80% ethanol twice before being allowed to dry for approximately 5 minutes. 12 µL of TE buffer 10/0.1 was then used to elute the DNA from the beads, vortexed briefly, and allowed to incubate at room temperature for 10 minutes prior to transfer of the supernatant into a new set of tubes The final concentration of DNA was determined using the DeNovix dsDNA High Sensitivity Kit. Libraries were sequenced either in single- or paired- end mode using Element AVITI platform with 150 cycles per end.

### ChIP-seq analyses

ChIP-seq analyses were performed similarly to how they were performed in previous studies^66^. In short, ChIP-seq reads had adapters trimmed using cutadapt^67^, using the standard Illumina TrueSeq adapters, with minimum length set to 15. Enrichment of reads on both pTarget as well as the BL21(DE3) genome was analyzed by mapping reads onto an indexed fasta file containing both a genome and plasmid sequence with two separate entries (one for the genome and one for the plasmid) utilizing bwa-mem2 and SAMtools packages^68,69^. BigWig files were created using deepTools2 bamCoverage command using the following parameters: -bs 1 -p 8 --normalizeUsing CPM --extendReads 200^70^. IGV was then used to visualize the mapped reads on both the pTarget plasmid as well as the whole genome^71^. Peak calling was done with MACS3 utilizing the following parameters: -g 4500000 --nomodel --extsize 400 -B -q 0.05 with respect to the non-immunoprecipitated input^72^. Sequence motifs were then identified using MEME with default parameters^73^.

### RFP fluorescence assay

pEffector and pTarget were double transformed into *E. coli* BL21(DE3). pEffector plasmids encoded IPTG inducible *arc* or Δ*arc* mutant genes. pTarget encoded either a transposon or *E. coli* promoter upstream of a synthetic strong ribosome binding site and a *mRFP1* gene. Single colonies were picked in biological replicates and grown in 1 mL of LB with antibiotics (to select for plasmids) for 8-10 hours for kinetic reads and 24-26 hours for endpoint reads in 96-well plates at 37 °C. For kinetic reads: After growth in the 96-well plate, 5 µL of liquid culture was transferred into 200 µL of LB, antibiotic, and either 0.0 or 0.5 mM IPTG into a 96-well optical bottom plate (Thermo Fisher Scientific). The plate was then placed into a Synergy Neo2 microplate reader (Biotek) for 13-16 hours with continuous shaking at 37 °C and parallel mRFP fluorescence signal and OD_600_ readings every 10 minutes. For endpoint reads: After growth in liquid cultures, a full 200 µL of liquid culture was aliquoted into a 96-well optical bottom plate (Thermo Fisher Scientific) and both mRFP fluorescence signal and OD_600_ were measured. mRFP signal was normalized to OD_600_.

### Plasmid interference assays

Plasmid interference assays were performed similarly to previous work^21^. pEffector plasmids encoding inducible *tnpB* genes were transformed into *E. coli* BL21(DE3), plated onto LB agar with spectinomycin (100 µg mL^−1^). Single colonies were selected to prepare chemically competent cells which were stored at −80 °C for future use. After thawing the competent cells on ice, 200 ng of each pTarget plasmid were transformed using heat shock at 42 °C for 45 seconds prior to recovery on ice for 8 minutes and recovery in 1 mL of LB per sample for 2 hours at 37 °C (while rotating). After recovery, cells were spun down for 5 minutes at 4000 g and the supernatant was discarded. The cell pellets were then resuspended in 50 µL LB and serially diluted (10) and plated onto LB agar containing spectinomycin (100 µg mL^−1^), kanamycin (50 µg mL^−1^), and 0.05 mM IPTG. These plates were grown for 16 hours at 37 °C overnight and imaged immediately after using a BioRad Gel Doc XR+ imager.

### TnpA excision assays

*E. coli* strain MG1655 cells were double-transformed with two plasmids. One of which was a mini-IS plasmid which contained both a kanamycin resistance gene, and the putative IS left and right ends surrounding a random 1000 bp sequence, forming a small artificial IS element. The second plasmid was one encoding an IPTG inducible *tnpA* gene as well as spectinomycin resistance. Both were utilized to double-transform into MG1655 cells and plated on LB-agar plates with spectinomycin (100 µg mL^−1^), kanamycin (50 µg mL^−1^), and 0.3 mM IPTG and grown overnight for 16 hours. Afterwards, colonies were scraped and resuspended in 200 µL of LB such that each sample contained about 3.2 10^8^ cells or an OD_600_ = 2.0 for 200 µL of LB. Cells were pelletted in a centrifuge at 4000 x*g* for 8 minutes prior to removal of the supernatant and resuspension in 80 µL of H2O. These cells were then incubated at 95 °C for 10 minutes and then the tubes were once again centrifuged at 4000 x*g* for 5 minutes and the supernatant/lysate moved into fresh tubes with the pellet discarded.

PCR reactions were performed using the flanking sequences adjacent to the mini-IS element designed to amplify the mini-IS element as well as the surrounding junction. PCR was performed utilizing 1 OneTaq Master Mix, 0.2 µM of each primer and 1 µL of the previously isolated lysate with a total PCR reaction volume of 20 µL. Reactions utilized an initial denaturation step at 94 °C for 30 seconds, followed by 30 cycles of denaturation at 94 °C for 15 seconds, annealing at 48 °C for 15 seconds, and then extension at 68 °C for 1 minute, and a final extension step at 68 °C for 5 minutes. Band sizes were resolved with gel electrophoresis on 1.5% agarose gel stained with SYBR Safe (Thermo Fisher Scientific). Bands of a length indicating a possible excision were extracted from the gel, column purified (Qiagen) and then were sequenced with Sanger sequencing (GENEWIZ).

### Data availability

Chromatin immunoprecipitation data will be made available through the Gene Expression Omnibus (GSE344792) at the time of publication. Datasets generated and analyzed in the current study are available from the corresponding authors on reasonable request.

### Code availability

No custom scripts were utilized for sequencing data analysis. Custom scripts used for bioinformatics are available upon request.

## ACKNOWLEDGMENTS

We thank T.M. Smith, A.J. Robinson, A.I. Palmieri, and R. Rafat for laboratory support; F.T. Hoffmann for guidance on ChIP-seq data processing; H.C. Le for advice on initial bioinformatic searches; and the JP Sulzberger Columbia Genome Center for next-generation sequencing support. S.H.S. was supported by NSF Faculty Early Career Development Program (CAREER) Award 2239685, a Pew Biomedical Scholarship, an Irma T. Hirschl Career Scientist Award, the Howard Hughes Medical Institute Investigator Program, and a generous startup package from the Columbia University Irving Medical Center Dean’s Office and the Vagelos Precision Medicine Fund.

## AUTHOR CONTRIBUTIONS

R.Z., Y.D.M., and S.H.S. conceived the study. Y.D.M. performed most experiments and initial bioinformatic analyses. T.W. performed the remaining bioinformatic analyses, including phylogenetic and gene association analyses. R.Z. mentored Y.D.M. and performed additional reporter assays. Y.D.M., R.Z., and S.H.S. wrote the manuscript, with input from all authors.

## COMPETING INTERESTS

S.H.S. is a co-founder and scientific advisor to Dahlia Biosciences, a scientific advisor to CrisprBits and Prime Medicine, and an equity holder in Dahlia Biosciences and CrisprBits.

Correspondence and requests for materials should be addressed to S.H.S.

## SUPPLEMENTARY TABLES

**Supplementary Table 1** | Arc homologs in Fig. S1 phylogenetic tree.

**Supplementary Table 2** | List of Arc homologs tested in this study.

**Supplementary Table 3** | Strains used in this study.

**Supplementary Table 4** | Description and sequence of plasmids used in this study.

**Supplementary Table 5** | Probes and oligonucleotides used in this study.

## REFERENCES

1. Haudiquet, M., de Sousa, J. M., Touchon, M. & Rocha, E. P. C. Selfish, promiscuous and sometimes useful: how mobile genetic elements drive horizontal gene transfer in microbial populations. Phil. Trans. R. Soc. B 377, 20210234 (2022).

2. Levin, H. L. & Moran, J. V. Dynamic interactions between transposable elements and their hosts. Nat Rev Genet 12, 615–627 (2011).

3. Pál, C. & Papp, B. From passengers to drivers: Impact of bacterial transposable elements on evolvability. Mobile Genetic Elements 3, e23617 (2013).

4. Cosby, R. L., Chang, N.-C. & Feschotte, C. Host–transposon interactions: conflict, cooperation, and cooption. Genes Dev 33, 1098–1116 (2019).

5. Sastre-Dominguez, J. et al. Plasmids promote antimicrobial resistance through insertion sequence-mediated gene inactivation. Nat Microbiol 11, 976–992 (2026).

6. Makarova, K. S. et al. Evolutionary classification of CRISPR–Cas systems: a burst of class 2 and derived variants. Nat Rev Microbiol 18, 67–83 (2020).

7. Kapitonov, V. V., Makarova, K. S. & Koonin, E. V. ISC, a Novel Group of Bacterial and Archaeal DNA Transposons That Encode Cas9 Homologs. Journal of Bacteriology 198, 797–807 (2016).

8. Altae-Tran, H. et al. The widespread IS200/IS605 transposon family encodes diverse programmable RNA-guided endonucleases. Science 374, 57–65 (2021).

9. Harmer, C. J., Moran, R. A. & Hall, R. M. Movement of IS26-Associated Antibiotic Resistance Genes Occurs via a Translocatable Unit That Includes a Single IS26 and Preferentially Inserts Adjacent to Another IS26. mBio 5, 10.1128/mbio.01801-14 (2014).

10. Dugan, J., Andersen, A. A. & Rockey, D. D. Functional characterization of IScs605, an insertion element carried by tetracycline-resistant Chlamydia suis. Microbiology 153, 71–79 (2007).

11. Elena, S. F., Ekunwe, L., Hajela, N., Oden, S. A. & Lenski, R. E. Distribution of fitness effects caused by random insertion mutations in Escherichia coli. Genetica 102, 349–358 (1998).

12. Reznikoff, W. S. Tn5 as a model for understanding DNA transposition. Molecular Microbiology 47, 1199–1206 (2003).

13. Tempel, S., Bedo, J. & Talla, E. From a large-scale genomic analysis of insertion sequences to insights into their regulatory roles in prokaryotes. BMC Genomics 23, 451 (2022).

14. Siguier, P., Gourbeyre, E., Varani, A., Ton-Hoang, B. & Chandler, M. Everyman’s Guide to Bacterial Insertion Sequences. Microbiol Spectr 3, MDNA3-0030–2014 (2015).

15. Toleman, M. A., Bennett, P. M. & Walsh, T. R. ISCR Elements: Novel Gene-Capturing Systems of the 21st Century? Microbiol Mol Biol Rev 70, 296–316 (2006).

16. Pasternak, C. et al. ISDra2 transposition in Deinococcus radiodurans is downregulated by TnpB. Molecular Microbiology 88, 443–455 (2013).

17. He, S. et al. IS200/IS605 family single-strand transposition: mechanism of IS608 strand transfer. Nucleic Acids Res 41, 3302–3313 (2013).

18. Chandler, M. et al. Breaking and joining single-stranded DNA: the HUH endonuclease superfamily. Nat Rev Microbiol 11, 525–538 (2013).

19. He, S. et al. The IS200/IS605 Family and ‘Peel and Paste’ Single-strand Transposition Mechanism. Microbiol Spectr 3, (2015).

20. Karvelis, T. et al. Transposon-associated TnpB is a programmable RNA-guided DNA endonuclease. Nature 599, 692–696 (2021).

21. Meers, C. et al. Transposon-encoded nucleases use guide RNAs to promote their selfish spread. Nature 622, 863–871 (2023).

22. Žedaveinytė, R., et al. Antagonistic conflict between transposon-encoded introns and guide RNAs. Science 385, eadm8189 (2024).

23. Su, C.-H. & Greene, R. C. Regulation of Methionine Biosynthesis in Escherichia coli: Mapping of the metJ Locus and Properties of a metJ+/metJ-Diploid. Proc Natl Acad Sci U S A 68, 367–371 (1971).

24. Weihofen, W. A., Cicek, A., Pratto, F., Alonso, J. C. & Saenger, W. Structures of ω repressors bound to direct and inverted DNA repeats explain modulation of transcription. Nucleic Acids Res 34, 1450–1458 (2006).

25. Schreiter, E. R. & Drennan, C. L. Ribbon-helix-helix transcription factors: variations on a theme. Nat Rev Microbiol 5, 710–720 (2007).

26. Baynham, P. J., Brown, A. L., Hall, L. L. & Wozniak, D. J. Pseudomonas aeruginosa AlgZ, a ribbon-helix-helix DNA-binding protein, is essential for alginate synthesis and algD transcriptional activation. Mol Microbiol 33, 1069–1080 (1999).

27. Baynham, P. J. & Wozniak, D. J. Identification and characterization of AlgZ, an AlgT-dependent DNA-binding protein required for Pseudomonas aeruginosa algD transcription. Molecular Microbiology 22, 97–108 (1996).

28. Waligora, E. A. et al. AmrZ Beta-Sheet Residues Are Essential for DNA Binding and Transcriptional Control of Pseudomonas aeruginosa Virulence Genes. Journal of Bacteriology 192, 5390–5401 (2010).

29. Li, X., Lozano-Madueño, C., Martínez-Alvarez, L. & Peng, X. A clade of RHH proteins ubiquitous in Sulfolobales and their viruses regulates cell cycle progression. Nucleic Acids Res 51, 1724–1739 (2023).

30. Oberer, M., Zangger, K., Gruber, K. & Keller, W. The solution structure of ParD, the antidote of the ParDE toxin–antitoxin module, provides the structural basis for DNA and toxin binding. Protein Sci 16, 1676–1688 (2007).

31. Bøggild, A. et al. The Crystal Structure of the Intact E. coli RelBE Toxin-Antitoxin Complex Provides the Structural Basis for Conditional Cooperativity. Structure 20, 1641–1648 (2012).

32. Kirsch, J. M. et al. Targeted IS-element sequencing uncovers transposition dynamics during selective pressure in enterococci. PLOS Pathogens 19, e1011424 (2023).

33. Lewis, L. A. et al. The left end of IS2: a compromise between transpositional activity and an essential promoter function that regulates the transposition pathway. J Bacteriol 186, 858–865 (2004).

34. Duval-Valentin, G., Normand, C., Khemici, V., Marty, B. & Chandler, M. Transient promoter formation: a new feedback mechanism for regulation of IS911 transposition. EMBO J 20, 5802–5811 (2001).

35. Dalrymple, B. & Arber, W. Promotion of RNA transcription on the insertion element IS30 of E. coli K12. EMBO J 4, 2687–2693 (1985).

36. Simons, R. W. & Kleckner, N. Translational control of IS10 transposition. Cell 34, 683–691 (1983).

37. Nagy, Z. & Chandler, M. Regulation of transposition in bacteria. Research in Microbiology 155, 387–398 (2004).

38. Camacho, C. et al. BLAST+: architecture and applications. BMC Bioinformatics 10, 421 (2009).

39. van Kempen, M. et al. Fast and accurate protein structure search with Foldseek. Nat Biotechnol 42, 243–246 (2024).

40. Altschul, S. F. et al. Gapped BLAST and PSI-BLAST: a new generation of protein database search programs. Nucleic Acids Res 25, 3389–3402 (1997).

41. Mears, K. S. et al. RNA-guided nucleases enable a gene drive of insertion sequences in plasmids. 2025.02.20.638934 Preprint at 10.1101/2025.02.20.638934 (2025).

42. Wiegand, T. et al. TnpB homologues exapted from transposons are RNA-guided transcription factors. Nature 631, 439–448 (2024).

43. Altae-Tran, H. et al. Diversity, evolution, and classification of the RNA-guided nucleases TnpB and Cas12. Proceedings of the National Academy of Sciences 120, e2308224120 (2023).

44. Solovyev, V. V. Solovyev, A Salamov (2011) Automatic Annotation of Microbial Genomes and Metagenomic Sequences. In Metagenomics and its Applications in Agriculture, Biomedicine and Environmental Studies (Ed. R.W. Li), Nova Science Publishers, p.61–78. in 61–78 (2011).

45. Zhang, X. et al. Petroclostridium xylanilyticum gen. nov., sp. nov., a xylan-degrading bacterium isolated from an oilfield, and reclassification of clostridial cluster III members into four novel genera in a new Hungateiclostridiaceae fam. nov. International Journal of Systematic and Evolutionary Microbiology 68, 3197–3211 (2018).

46. Meng, E. C. et al. UCSF ChimeraX: Tools for structure building and analysis. Protein Science 32, e4792 (2023).

47. Abramson, J. et al. Accurate structure prediction of biomolecular interactions with Alpha-Fold 3. Nature 630, 493–500 (2024).

48. Pei, J. & Grishin, N. V. AL2CO: calculation of positional conservation in a protein sequence alignment. Bioinformatics 17, 700–712 (2001).

49. Ziemann, M. et al. CvkR is a MerR-type transcriptional repressor of class 2 type V-K CRISPR-associated transposase systems. Nat Commun 14, 924 (2023).

50. Ziemann, M. et al. Analysis of tracrRNAs reveals subgroup V2 of type V-K CAST systems. Microlife 6, uqaf020 (2025).

51. Fauconnier, A. et al. Dual regulatory role of IS91-encoded Orf121 in IS91 transposition. Commun Biol 9, 667 (2026).

52. Mohsen, J. J., Martel, A. A. & Slavoff, S. A. Microproteins—Discovery, structure, and function. PROTEOMICS 23, 2100211 (2023).

53. Fesenko, I. et al. The hidden bacterial microproteome. Molecular Cell 85, 1024–1041.e6 (2025).

54. Sberro, H. et al. Large-Scale Analyses of Human Microbiomes Reveal Thousands of Small, Novel Genes. Cell 178, 1245–1259.e14 (2019).

55. Fremin, B. J. et al. Thousands of small, novel genes predicted in global phage genomes. Cell Reports 39, 110984 (2022).

56. Altschul, S. F., Gish, W., Miller, W., Myers, E. W. & Lipman, D. J. Basic local alignment search tool. Journal of Molecular Biology 215, 403–410 (1990).

57. Eddy, S. R. Accelerated Profile HMM Searches. PLOS Computational Biology 7, e1002195 (2011).

58. Mistry, J. et al. Pfam: The protein families database in 2021. Nucleic Acids Res 49, D412–D419 (2021).

59. Xie, Z. & Tang, H. ISEScan: automated identification of insertion sequence elements in prokaryotic genomes. Bioinformatics 33, 3340–3347 (2017).

60. Cantalapiedra, C. P., Hernández-Plaza, A., Letunic, I., Bork, P. & Huerta-Cepas, J. egg-NOG-mapper v2: Functional Annotation, Orthology Assignments, and Domain Prediction at the Metagenomic Scale. Mol Biol Evol 38, 5825–5829 (2021).

61. Steinegger, M. & Söding, J. MMseqs2 enables sensitive protein sequence searching for the analysis of massive data sets. Nat Biotechnol 35, 1026–1028 (2017).

62. Katoh, K., Misawa, K., Kuma, K. & Miyata, T. MAFFT: a novel method for rapid multiple sequence alignment based on fast Fourier transform. Nucleic Acids Res 30, 3059–3066 (2002).

63. Capella-Gutiérrez, S., Silla-Martínez, J. M. & Gabaldón, T. trimAl: a tool for automated alignment trimming in large-scale phylogenetic analyses. Bioinformatics 25, 1972–1973 (2009).

64. Price, M. N., Dehal, P. S. & Arkin, A. P. FastTree 2 – Approximately Maximum-Likelihood Trees for Large Alignments. PLOS ONE 5, e9490 (2010).

65. Bonocora, R. P. & Wade, J. T. ChIP-Seq for Genome-Scale Analysis of Bacterial DNA-Binding Proteins. in Bacterial Transcriptional Control: Methods and Protocols (eds Artsimovitch, I. & Santangelo, T. J.) 327–340 (Springer, New York, NY, 2015). doi:10.1007/978-1-4939-2392-2_20.

66. Hoffmann, F. T. et al. Selective TnsC recruitment enhances the fidelity of RNA-guided transposition. Nature 609, 384–393 (2022).

67. Martin, M. Cutadapt removes adapter sequences from high-throughput sequencing reads. EMBnet.journal 17, 10–12 (2011).

68. Li, H. et al. The Sequence Alignment/Map format and SAMtools. Bioinformatics 25, 2078–2079 (2009).

69. Vasimuddin, Md., Misra, S., Li, H. & Aluru, S. Efficient Architecture-Aware Acceleration of BWA-MEM for Multicore Systems. in 2019 IEEE International Parallel and Distributed Processing Symposium (IPDPS) 314–324 (2019). doi:10.1109/IPDPS.2019.00041.

70. Ramírez, F., Dündar, F., Diehl, S., Grüning, B. A. & Manke, T. deepTools: a flexible platform for exploring deep-sequencing data. Nucleic Acids Res 42, W187–W191 (2014).

71. Robinson, J. T. et al. Integrative genomics viewer. Nat Biotechnol 29, 24–26 (2011).

72. Zhang, Y. et al. Model-based Analysis of ChIP-Seq (MACS). Genome Biol 9, R137 (2008).

73. Bailey, T. L. et al. MEME Suite: tools for motif discovery and searching. Nucleic Acids Res 37, W202–W208 (2009).

