## Supplementary Figures for "Arc represses gene expression in IS*605*-family transposons"

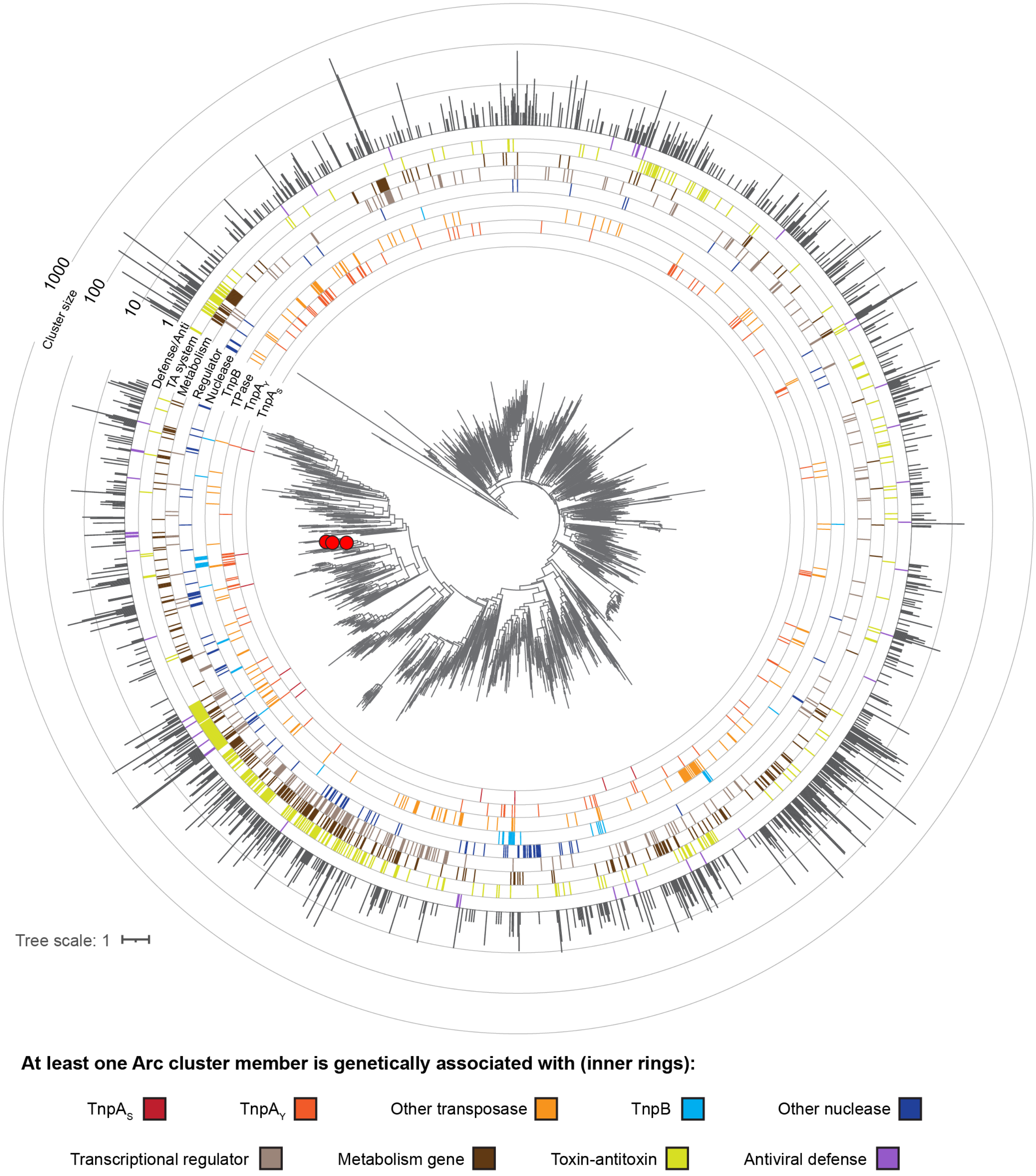


**Figure S1 | Phylogenetic tree of Arc proteins with labeled gene associations.** Phylogenetic tree constructed from 2,175 Arc cluster representatives. Inner rings visualize proteins encoded immediately proximal to *arc*, for at least one member of the cluster. The outer ring depicts the number of sequences in each cluster. Arc-like RHH homologs are associated with functionally diverse genes, including metabolic components and toxin-antitoxins. Cluster members and representatives are listed in **Supplementary Table 1**. This tree of cluster representatives is an approximate ML tree used to order representatives for visualization, and deep relationships should not be interpreted due to the high sequence divergence across this dataset of Arc homologs.

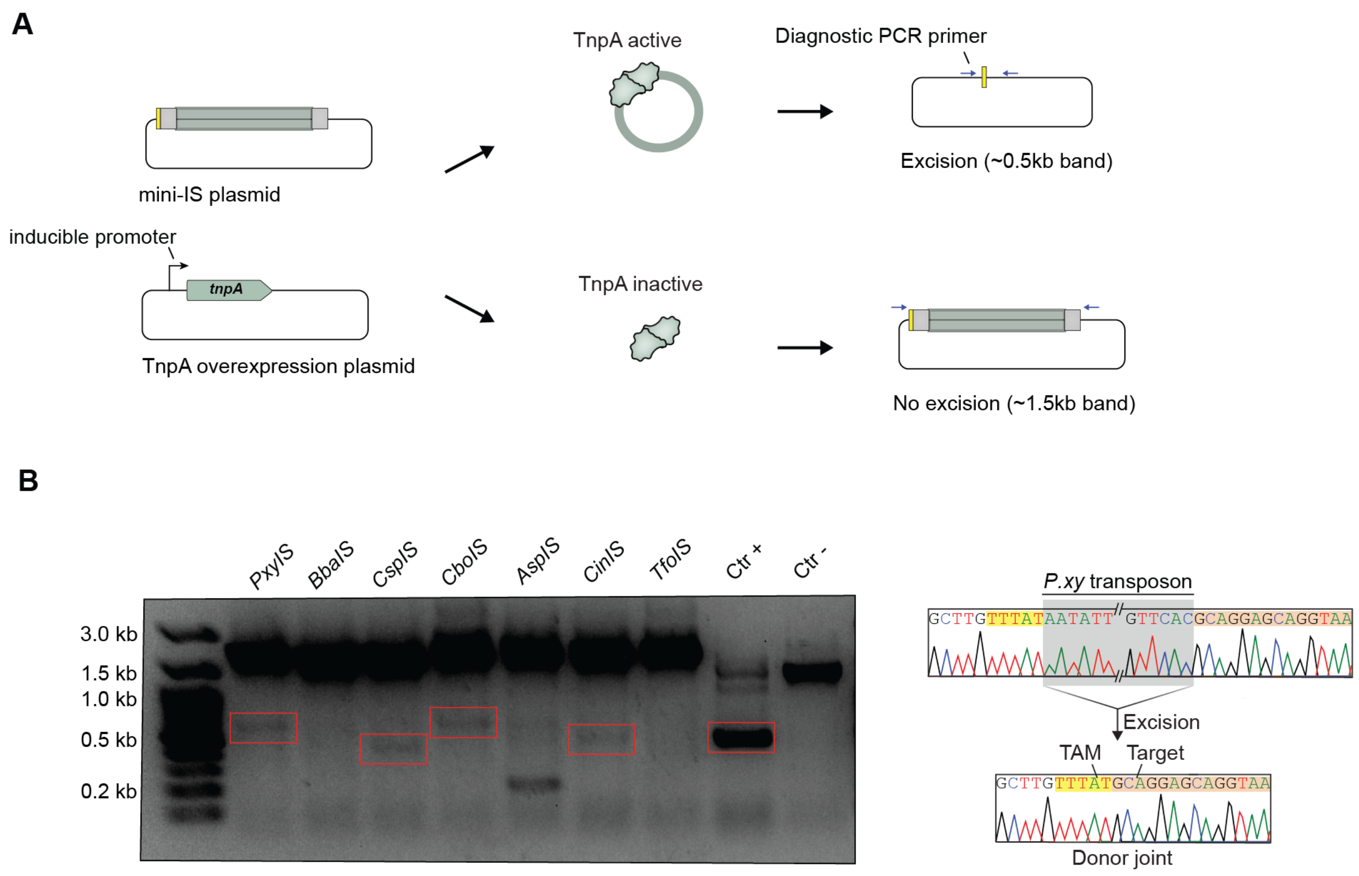


**Figure S2 | Excision PCR assays demonstrate TnpA excision activity**

**A)** Schematic demonstrating PCR assays to check for TnpA activity. The assay included a mini-IS plasmid containing a 1kb stuffer sequence with the native left and right transposon ends and a TnpA overexpression plasmid encoding the corresponding TnpA encoding gene with inducible expression. Upon double transformation and induction, active TnpAs would excise the mini-IS from some proportion of the mini-IS plasmids. By using PCR primers to PCR the junction we were able to measure the excision activity of TnpA for seven systems.

**B)** PCR bands for seven autonomous Arc-containing IS605 elements alongside a positive control TnpA known for strong excision activity and a negative control dTnpA which is excision inactive. *P. xylanilyticum* as well as a number of other systems show excision activity. Sequencing revealed excision junctions for most bands besides the band for *A. sp* which formed due to mispriming. On the right hand side is an example Sanger sequencing output in which the plasmid sequence includes the *P. xy* mini-IS transposon whereas the excision band sequence is absent of this transposon sequence.

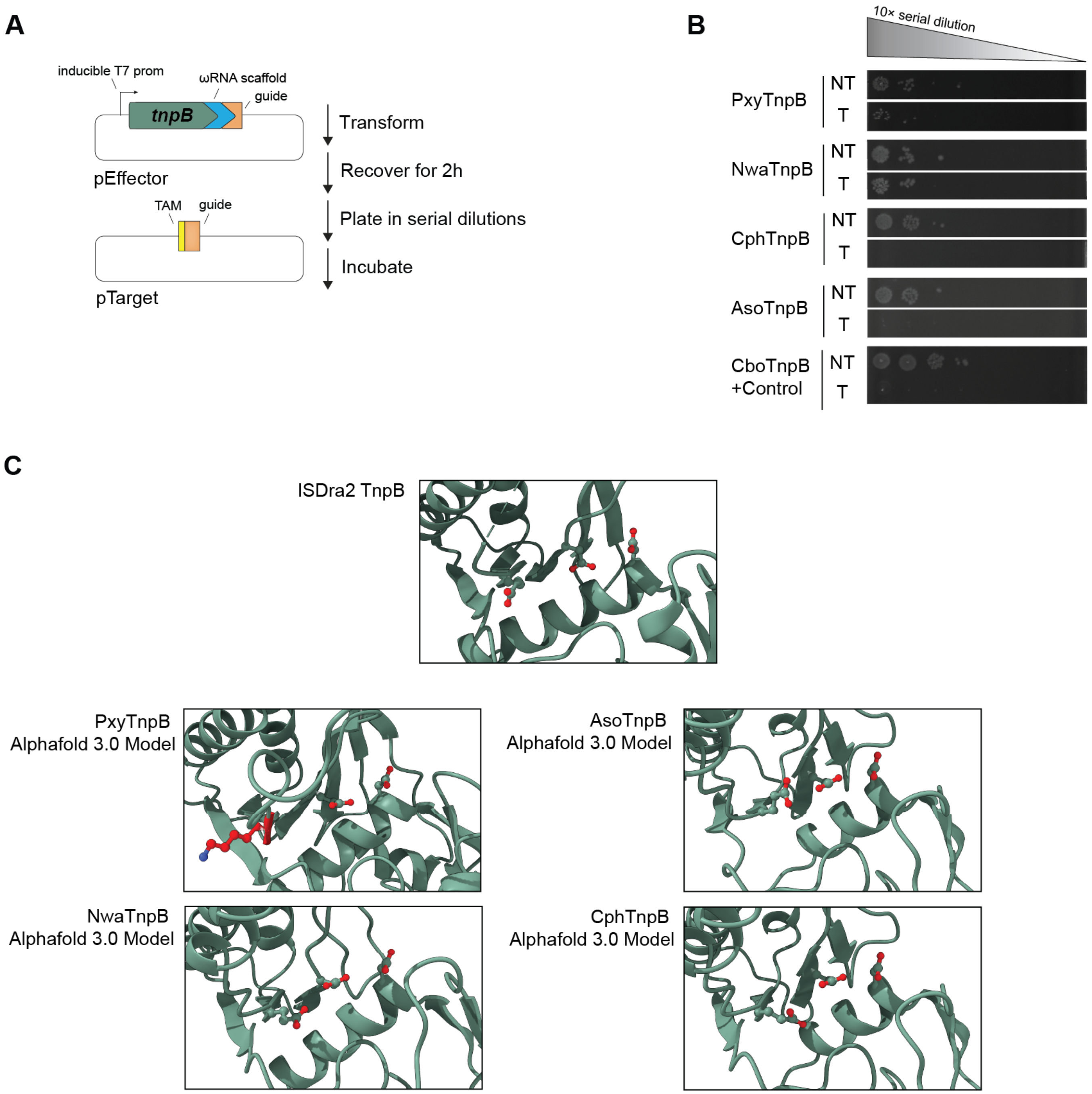


**Figure S3 | Functionality of Arc associated TnpB endonucleases.**

**A)** Constructs and steps involved in TnpB cleaving spot assays. The assay involved the expression of TnpB with a guide RNA sequence designed to target a plasmid with a kanamycin resistance gene. Upon cleavage by active TnpB, kanamycin resistance would be lost leading to cell death when plated on LB agar with kanamycin present

**B)** Results of TnpB cleaving spot assays. NT refers to a non-targetting guide control and T refers to a guide that targets pTarget as shown in Fig. S6a above. We see that *Aneurinibacillus soli* and Clostridium Phage D-1873's Arc associated TnpBs appear to be active as observed by the clearance of colonies in the targeting condition. In contrast, it appears *Nitrosococcus wardiae* and *Petroclostridium xylanilyticum* have little to no TnpB cleaving activity.

**C)** Alphafold 3.0 models of tested TnpB homologs. Above is a reference structure for ISDra2 TnpB, a known, functional IS605 TnpB. Below are Alphafold 3.0 models of all the tested Arc-associated TnpBs. Most show an intact DDE catalytic motif with minimal perturbation to the canonical TnpB structure, but notably *P. xylanilyticum* has lost the key glutamate residue necessary for TnpB activity.

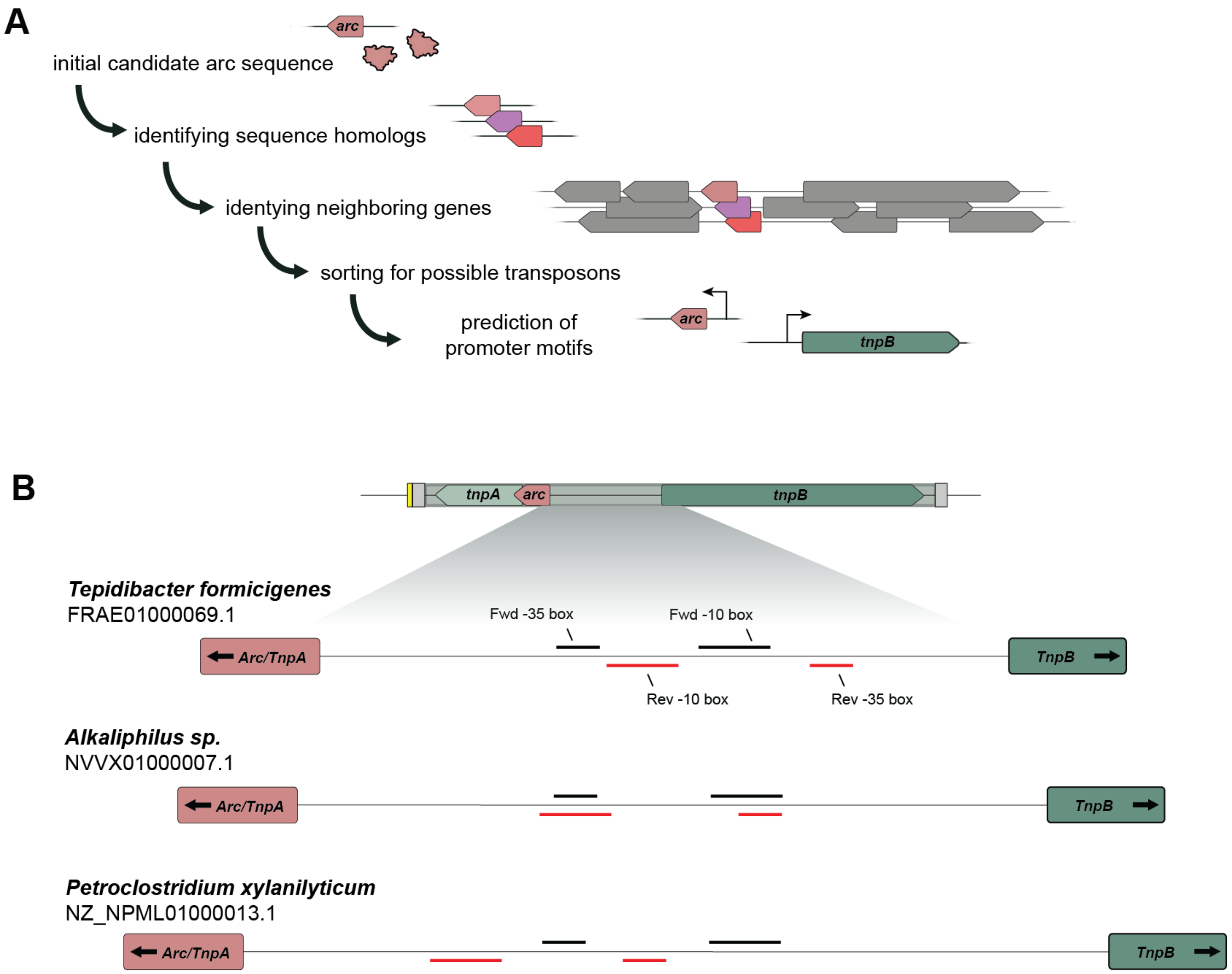


**Figure S4 | Bioinformatic identification of promoters for Arc containing autonomous IS605 elements**

**A)** Schematic demonstrating the identification of arc-associated IS605 transposons and potential promoters regulating gene expression for further testing. The process involved first identifying homologs to already known Arc proteins encoded within IS605 transposons, then using neighboring TnpA and TnpB genes to infer possible transposon associations. By looking for closely adjacent TnpA, TnpB, and Arc genes, as well as the existence of replicate sequences in the genome, implying possibly active transposition, a number of possible transposon sequences were found. Possible promoter motifs were then predicted utilizing BPROM.

**B)** Example transposon promoter organizations. All autonomous transposons tested encoded TnpA, TnpB, and Arc encoded as shown in the diagram above, with TnpB encoded in one direction and TnpA and Arc encoded in the reverse direction. Inputting the intergenic sequence of each transposon often resulted in one promoter predicted in the TnpB orientation and one promoter predicted in the Arc/TnpA orientation. These motifs had variable spacing between different transposon homologs. Above in black are the predicted forward promoter motifs, and below in red are the predicted reverse promoter motifs for three representative species each with an Arc-containing IS605 element.

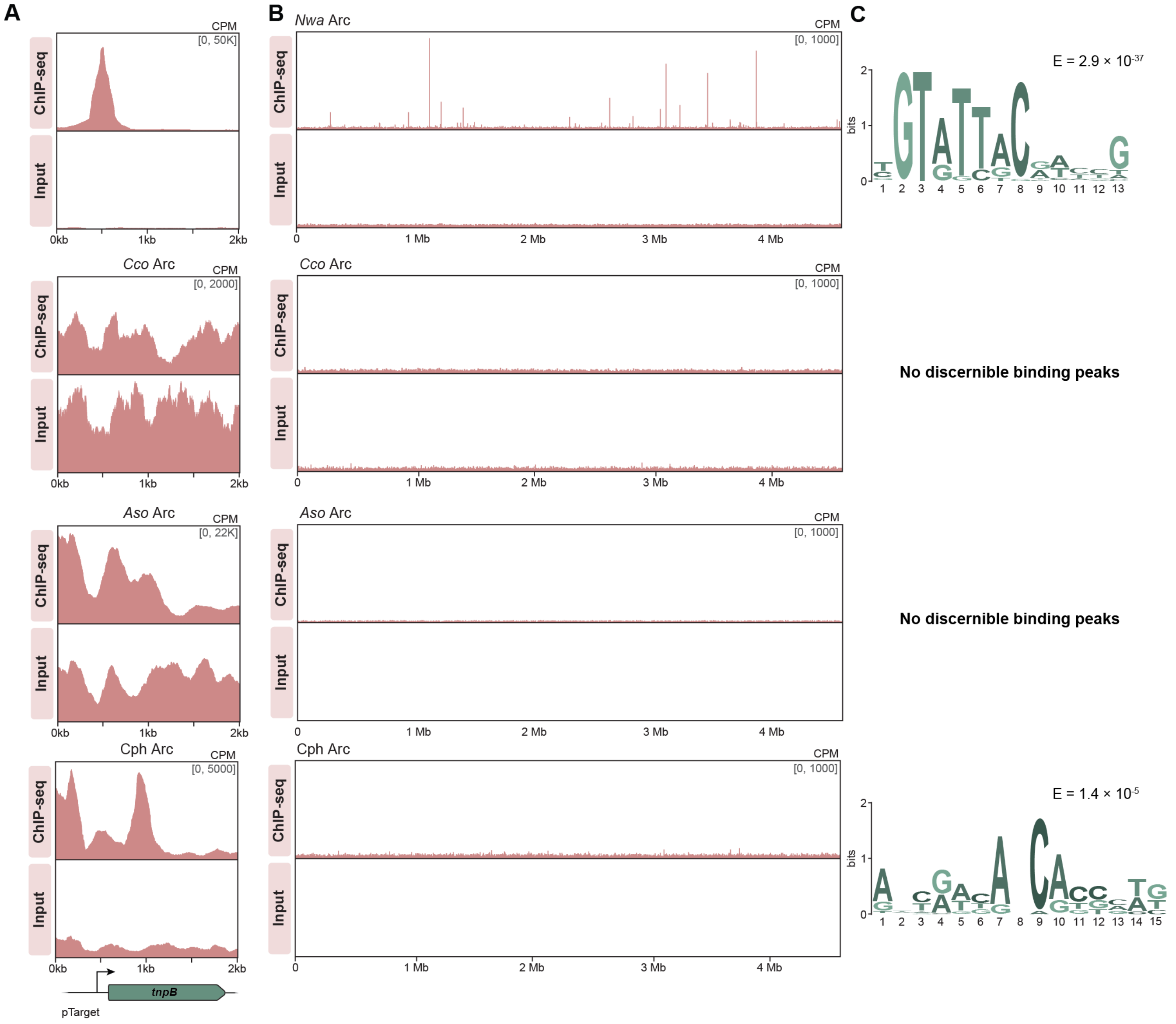


**Figure S5 | Arc binding is not always present in all Arc containing IS605 elements.**

**A)** ChIP-seq transposon binding data for experiments done with *N. wardiae*, *Candidatus competibacteraceae*, *A. soli*, and Clostridium phage D-1873 derived Arcs. While some Arc proteins show binding peaks near the transposon's promoter or TnpB coding region, others show no evidence of binding to the transposon sequence at all.

**B)** ChIP-seq data mapped to the genome of BL21(DE3) *E. coli*. In transposons that Arc failed to bind to, we see no other enrichment over any genomic DNA, implying that these Arcs may fail to bind DNA in any sequence specific manner.

**C)** Binding motifs for arc-like RHH repressors for *N. wardiae* and Clost. phage D-1873 derived from the genomic peaks using MEME-ChIP to determine consensus sequences. *A. so* and *C. co* show no consensus peaks due to no notable peaks being found within the ChIP-seq data, further supporting that these Arcs are not binding DNA in any specific manner. E, E-value significance.

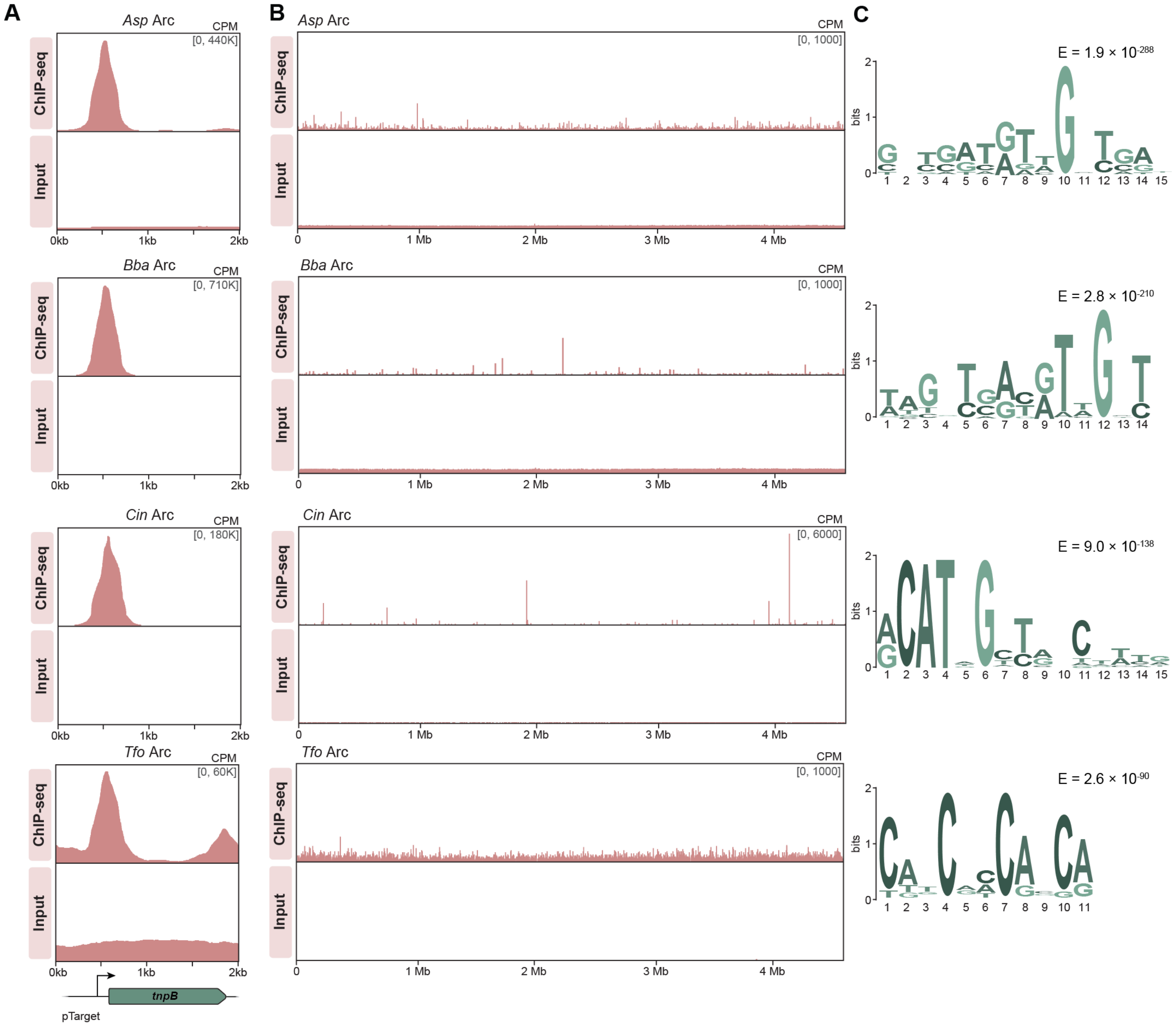


**Figure S6 | ChIP-seq analysis shows multiple Arc homologs bind back to transposon sequence with unique consensus sequences.**

**A)** ChIP-seq transposon binding data for experiments done with *Alkaliphilus sp, Crassaminicella indica, Tepidibacter formicigenes,* and a Bacilli bacterium. All these systems contain IS605 elements encoding Arc, and upon overexpression of Arc in the presence of transposon sequence we see strong enrichment over the transposon promoter for all systems confirming that other homologs also bind back to the transposon sequence, likely also playing an autoregulatory function.

**B)** ChIP-seq data mapped to the genome of BL21(DE3) *E. coli*. Similar to other binding active Arc proteins, enrichment of reads in the genome is substantially less than the enrichment over the transposon's native promoter sequence, implying minimal binding events to partial motifs.

**C)** Binding consensus motifs for Arcs determined from genomic peaks using MEME-ChIP. E, E-value significance.
